# Occurrence of snail intermediate host of schistosomiasis: implications for schistosomiasis control based on mass drug administration of praziquantel in Benue State, Nigeria

**DOI:** 10.64898/2026.01.22.701117

**Authors:** James Agada Okete, Eme Effiong Etta, Patricia Edodi Ikika, Theophilus Effiong Akpe, Benedict Onu Onoja

## Abstract

Schistosomiasis is a neglected disease that is still ravaging many rural community dwellers in Nigeria. Different species of freshwater snails, such as *Bulinus* and *Biomphalaria*, play passive roles in the transmission of schistosomiasis. A study on the occurrence of snail intermediate hosts of schistosomes was conducted following mass praziquantel administration in Katsina-Ala, Benue State, Nigeria. The objective was to determine the prevalence of freshwater snails. incriminated in transmitting schistosomiasis and their cercaria infectivity. A total of 24,812 freshwater snails belonging to six different genera (*Bulinus globosus, B. forskalii, Pila ovata, Lanistes lybicus, Biomphalaria pfeifferi, and Lymnaea natalensis* were encountered. Snails collected were kept under laboratory conditions for cercaria emergence. *B. globosus, B. forskalii*, and *B. pfeifferi* were identified as vectors of Schistosomes because they secreted cercaria of S. *haematobium* and *S. mansoni*, respectively. Parasite infection was higher in the ponds (83.33 %); followed by dams (68.57 %)) and streams (53.33 %). There was positive correlation (r = 0.657; P < 0.05) between the amount of rainfall and abundance of all snail species and their infection rate collected except *Biomphalaria* species that showed a negative correlation. The studies, therefore, call for Consideration of snails control to be integrated with Praziquantel Mass Drug Advocacy Campaigns so as to achieve transmission interruption.

## 1.0 Introduction

Schistosomiasi*s* is commonly called bilharziasis. It is among the principal public health diseases caused by parasitic trematodes; hence, its name, snail fever. According to Okete *et al*. (2022), this disease is prevalent in both tropical and subtropical countries. It is one of the “forgotten diseases of forgotten people,” even though it occurs in over 73 countries globally. Although it is of immense public health importance in humans, it has received considerable neglected attention in terms of control, maybe because it does not have a dramatic effect (Stafanie *et al*., 2016).

Based on prevalence, morbidity burden, and socio-economic and public health importance, it is ranked as the third parasitic disease in the world. Chigozie *et al*. (2009) also stated that more than 200 million people are infected worldwide, with 120 million symptomatic and 20 million with severe illness, with varying pathological presentations. An estimated 80% of the most severely affected individuals are now concentrated in Africa, and about 400 million more people are at risk of infection (Omudu and Iyough, 2017). According to Engels (1997), the diseases resulting from the species of schistosomes are categorized into two main forms with respect to their niches in the definitive hosts.

The transmission of *Schistosoma haematobium* is initiated by the shedding of cercariae by *Bulinu*s species, the favorable intermediate host. The genus *Bulinus* is found in the majority of African Countries and their adjoining regions, representing one of the most important components of freshwater zoobenthos (Almos,2024). Their shells provide a substrate for many organisms to create epibiotic communities of predominantly facultative nature (Tomasz, 2019). They are adapted to a wide range of ecological conditions and are found worldwide where there is sufficient water for their survival (Borong *et al*., 2020).

To encourage effective management of helminth diseases, including schistosomiasis, the 54th World Health Assembly (WHA) endorsed resolution 54.19 in 2001. Consequently, many sponsors, including NGOs and governments in endemic regions, are now paying keen interest to schistosomiasis and other neglected tropical diseases (NTDs) control. For instance, there was mass praziquantel treatment in Uganda and Tanzania in 2003 (Guyatt, 2010) and Nigeria in 2017 (Personal discussion with Prof. E. Braid). According to Bruno and Omar (2016), a cost-effective approach in the delivery of praziquantel through the schools was adopted in all these countries. However, elimination or reduction of the disease through this school-based method of treatment has been challenged by re-infection since other members of the community who may harbour the infection and potential sources of contamination or transmission are left out (Gower *et al*., 2007).

It is no doubt that schistosomiasis is still a public health menace in geographically restricted areas in Nigeria despite many years of extensive research. In endemic areas, it is a grave cause of morbidity and mortality. According to Stefanie *et al*. (2016), treatment with praziquantel (PZQ) is effective at reducing or eliminating active infection but does not prevent reinfection. The Neglected Tropical Diseases (NTD) Control unit of the Federal Ministry of Health (FMOH) has commenced schistosomiasis control and elimination programs based on one round of Praziquantel Mass Drug Administration (PMDA) targeted primarily at primary school age children in schools and out-of-schools in all the states of the federation, including Benue State as part of fulfilling the mandate of the 54th World Health Assembly Resolution 54.19 (Personal discussion with Professor E. Braid; Dr. Debam, Mark, NTD coordinator Benue State chapter). However, there is a paucity of firsthand information on the extent to which or whether or not the administration of praziquantel affects the distribution and abundance of freshwater snails, the intermediate host of schistosomiasis. It was based on this that an evaluation was carried out after mass praziquantel administration to determine the occurrence of freshwater snails and their potential for transmitting schistosomiasis.

## 2.0 Materials and Methods

### 2.1. Study Location

This study was carried out in Katsina-Ala Local Government Area of Benue State, Nigeria. Benue State is located in the Middle-Belt Region of Nigeria. The state lies within the lower river Benue trough in the middle-belt region of Nigeria. It lies between longitude 7.47 and 10.0 east of the Greenwich meridian and latitude 6.25 and 8.8 north of the equator. The state, which has a total landmass of 34,059 square kilometers, shares boundaries with five other states of the federation, namely: Nasarawa State to the North, Taraba State to the East, Cross River State to the South, Enugu State to the South-west and Kogi State to the West.

Benue State is an agrarian state; it is divided into three geopolitical zones, sometimes referred to as agricultural Zones, namely, Zone A, B, and C. The location of the experiment was in Zone A, which is Katsina-Ala. It is a cosmopolitan settlement on the North Bank of the river from which the town takes its name. The people of Katsina-Ala Local Government Area are predominantly farmers. Over 75% of the population engages in agriculture, making agriculture the mainstay of the economy of the people.

### 2.2 Study design

These studies employed a cross-sectional research method, which was conducted in across different water bodies (ponds, dams, and streams) in different communities where human water contact was eminent.

### 2.3 Ethical approval and informed consent

Ethical Clearance (Ref. No: MOA/STA/204/VOL.1/71) used during this study was granted by the Research and Ethics Committee unit, Benue State Ministry of Agriculture and Natural Resources. Verbal consent and permission to carry out a snail survey in water bodies were obtained from the different community heads in the Katsina-Ala Local Government Area based on an introductory letter and ethical clearance. The introduction letter was issued by my supervisors on behalf of the Departmental Research Ethics Committee.

### 2.5 Reconnaissance Survey

Before the commencement of the research, we took a familiarization tour of all the communities that were chosen to obtain their full consent. We met with the Heads of communities to present the reasons for doing the work and its potential benefits.

### 2.6 Praziquantel Administration

The Federal Ministry of Health provided the praziquantel through the Neglected Tropical Diseases (NTDs) Network in the State

Mass administration of the drug in a standard single dose of 40 mg/kg according to body weight to the respective schools/schoolchildren was done by the Local Government Area NTDs coordinator, assisted by some school teachers.

### 2.7 Malacological and transmission studies

To establish continued transmission and re-infection, freshwater bodies designated and classified into streams, ponds, and dams within the study areas were located with the aim of determining *the presence of Schistosoma* cercariae in freshwater snails. Criteria for selection of freshwater bodies for snail collection include proximity to human habitation and human water contact activities. Sampling stations were selected based on human contact points.

The water bodies were painstakingly examined at monthly intervals from April 2018 to March 2019 using a long scooping net constructed by the researcher as outlined by Hairston (1990) and Azim and Ayad (1984) and employed by Omudu and Iyough (2017) and Asor (1995). The scoop, made up of mosquito netting with a mesh size of 1 mm, was 30 cm deep and attached to a square wooden frame of 20 cm in diameter, nailed to a long wooden handle of 95 cm in length.

Snails collected were put in experimental containers (sterile plastic containers) containing damp and decaying leaves with water from the habitats (Okete *et al*, 2015) and carried to the postgraduate/Parasitological Research Laboratory, Department of Zoology, University of Agriculture, Makurdi. In the laboratory, snails were identified based on Principal Component Analysis and morphological features using field guide keys outlined by WHO (2009); Christensen and Frandsen (1985), and employed by Asor (1995). Snail infection rate was evaluated using the free emergence method.

### 2.8 Examination of snails for cercaria infectivity

Snails were screened for patent trematode infection using the cercaria shedding method in line with Christensen and Frandsen’s (1985) procedures, with a few modifications. Each was placed singly in 250 ml plastic containers (experimental container) of 7. 5cm depth x 13 cm length 8.5cm breadth containing 100 ml of water collected from the snails’ habitat and exposed to strong artificial illumination (60 W bulbs) for a period of 4-6 hours to influence cercaria shedding. Water from the experimental container was collected and centrifuged at 3000 rpm for 5 minutes. 2-3 drops of the aliquots were collected with a pipette and placed on a glass slide. Lugol iodine was added to the deposits /aliquots on the slide to immobilize and stain the cercariae if present. It was then examined by a light microscope (Olympus, German series) under X 10 and X 40 objectives. All brevifurcate cercariae were identified as those of *S. haematobium* by morphology as previously described by Christensen and Frandsen (1985). The number of each snail species was recorded, and the monthly infection rate per site was recorded.

### 2.9 Statistical Analysis

Chi-square (χ^2^) was used to test significant differences in the abundance of snails and their infection rate in different sampling locations, while Pearson’s coefficient of correlation was employed to determine snail infection and its relationship with rainfall.

## 3.0 Results

### 3.1 Abundance of freshwater snail species in freshwater habitats in Katsina-Ala, Benue State, Nigeria

Of the species of snails collected, *Bulinus globosus* was the highest (7,302) number collected, followed by *P. ovata* (5,268), *L. lybicus* (4,347), *B. pfeifferi* (3,601), *L. natalensis* (2,246), and *B. forskalii* (2, 048) (Table 1). The overall numerical abundance of all snail species collected from all habitats was 24,812. Monthly fluctuation of schistosome infection in *Bulinus globosus, Bulinus forskalii*, and *Biomphalaria* in different habitats in Katsina-Ala, Benue State, Nigeria.

**Tale 1:**
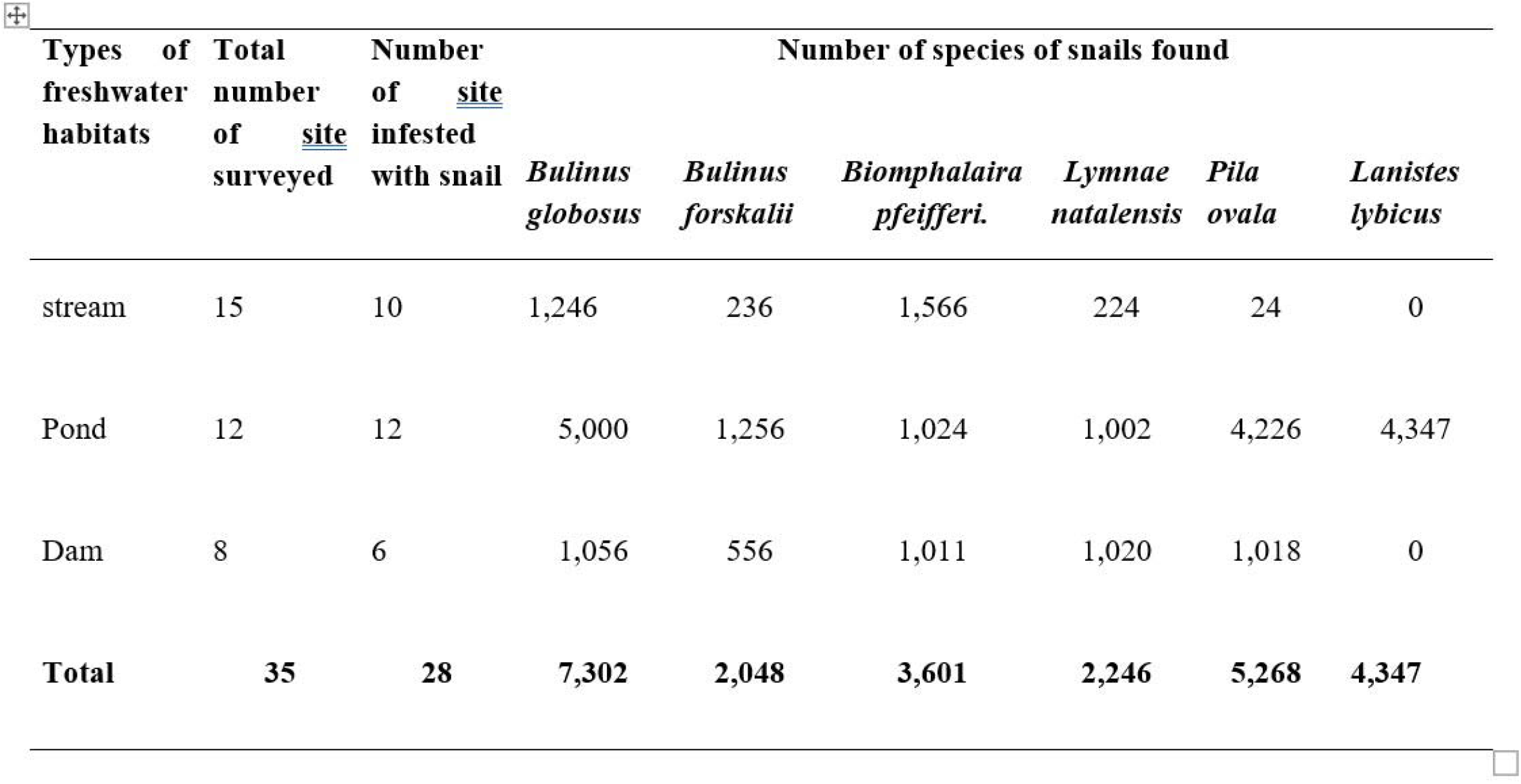
Abundance of freshwater snail species in freshwater habitats in Katsina-Ala, Benue State, Nigeria.

| Types of freshwater habitats | Total number of <u>site</u> surveyed | Number of <u>site</u> infested with snail | Number of species of snails found |  |  |  |  |  |
| --- | --- | --- | --- | --- | --- | --- | --- | --- |
|  |  |  | *Bulinus globosus* | *Bulinus forskalii* | *Biomphalaira pfeifferi.* | *Lymnae natalensis* | *Pila ovala* | *Lanistes lybicus* |
| stream | 15 | 10 | 1,246 | 236 | 1,566 | 224 | 24 | 0 |
| Pond | 12 | 12 | 5,000 | 1,256 | 1,024 | 1,002 | 4,226 | 4,347 |
| Dam | 8 | 6 | 1,056 | 556 | 1,011 | 1,020 | 1,018 | 0 |
| **Total** | **35** | **28** | **7,302** | **2,048** | **3,601** | **2,246** | **5,268** | **4,347** |

In stream habitats, infected *Bulinus globosus* occurred throughout the months in the rainy season, except in the dry season, where snail infection did not occur. A total of 520 snails were infected. However, in pond and dam habitats, infected *B. globosus* occurred in all the months (both dry and rainy season) with 3,074 snails infected in the pond and 840 snails infected in the dams. However, peak infection occurred during the rainy season (the month of September)-Figure 1. There was a positive correlation between infected snails and the amount of rainfall in all the habitat types.

**FIG 1:**
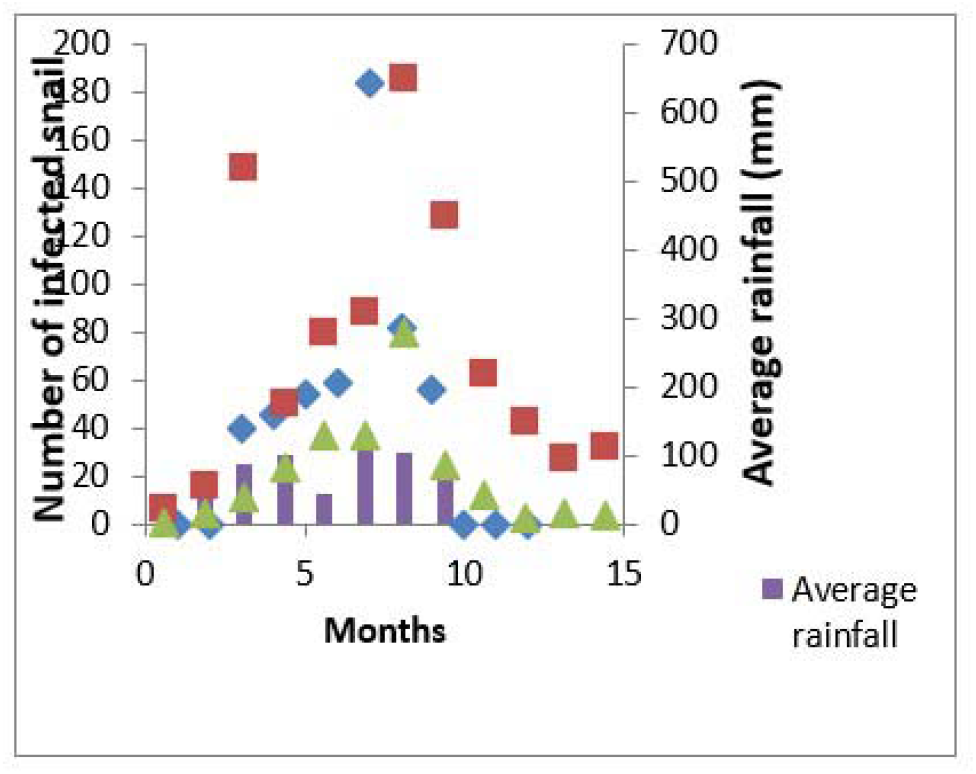
Fluctuation in numbers of infected *B. globosus* snails in stream, ponds, and dams, and their relationship to rainfall

Infected *B. forkalii* did not occur in stream habitats in all the months except in September, October, and November, with a total of 13 infected snails. In ponds and dams’ habitats, infected *B. forkalii* occurred in all the months except in February and March, with a total of 720 infected snails collected in ponds and 487 infected snails collected in dams (Figure 2). Peak infection of snails occurred more in the rainy season than in the dry season. There was a positive (r = 0.057; P < 0.05) correlation between infected *B. forkalii in* all the habitat types except in stream habitats, where there was a negative correlation (r = −0.023; P > 0.05).

**FIG 2:**
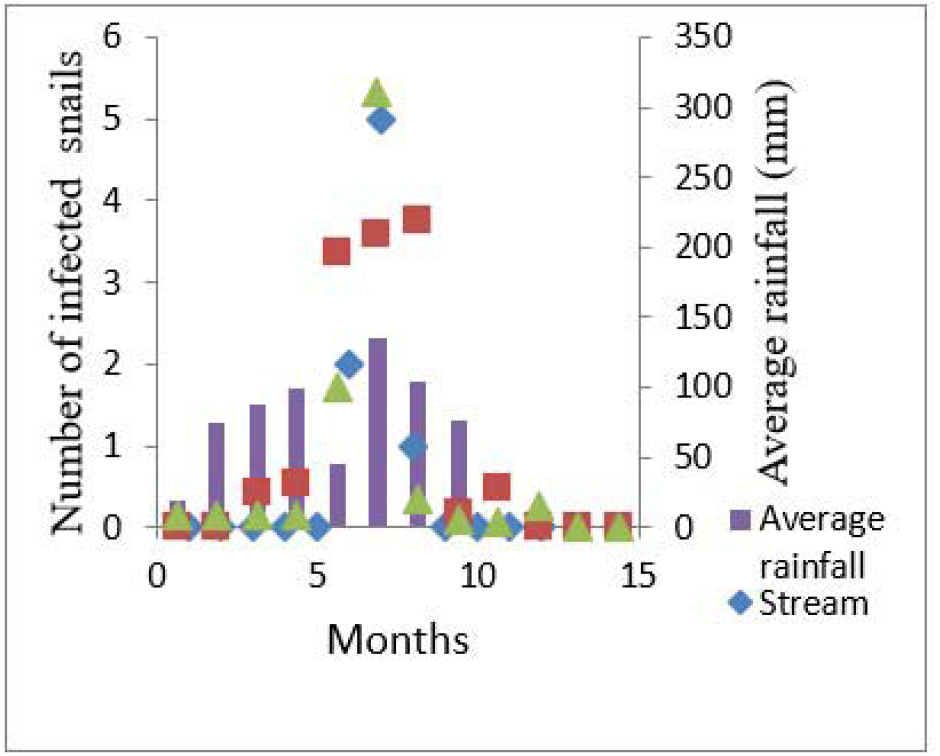
Fluctuation in numbers of infected *B. forskalii* snails in stream, ponds, and dams, and their relationship to rainfall

**FIG 3:**
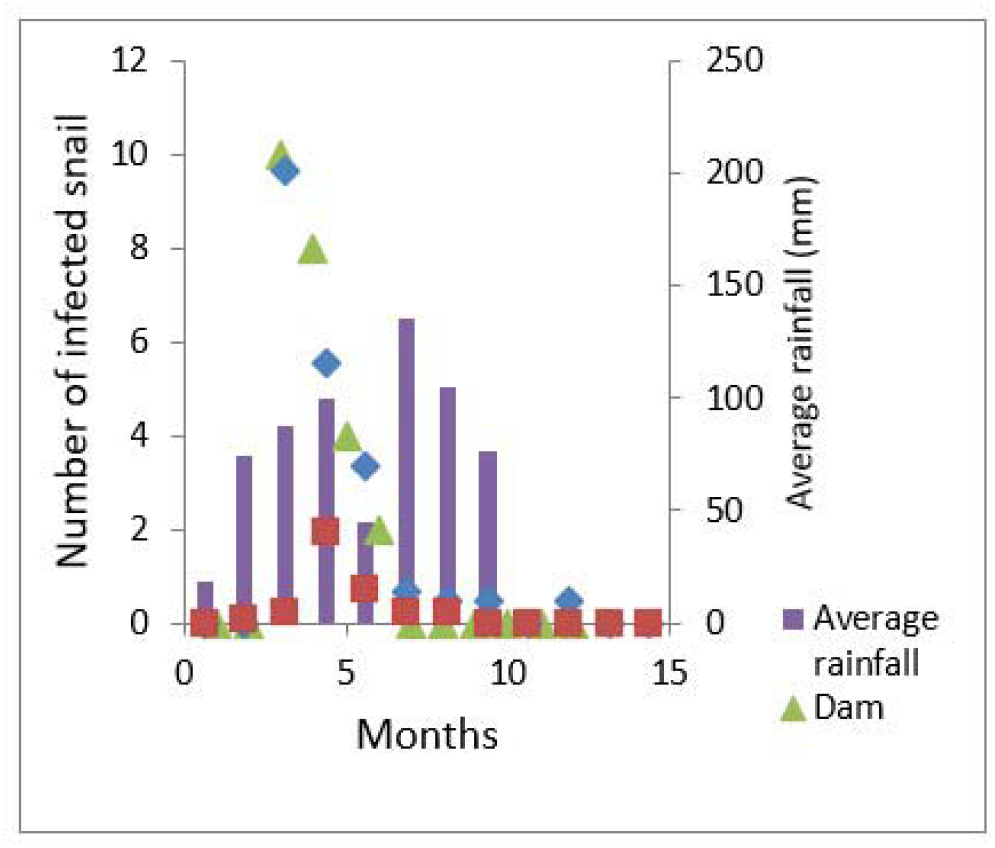
Fluctuation in numbers of infected *B. pfeifferi* snails in stream, ponds, and dams, and their relationship to rainfall

Infected *B. pfeifferi* occur in all the months except in April, May, December, January, and March in stream habitats with 428 infected snails. In ponds and dams’ habitats, infected *B. pfeifferi* occurred in all the months except in October-March, with a total of 72 infected snails collected in ponds and 24 infected snails collected in dams (Figure 2). Peak infection of snails occurred more in the month of June. There was a negative correlation (r = −0.023; P > 0.05) between infected *B. pfeifferi* and rainfall in all the habitat types.

## 4.0 Discussion

### (i) Occurrence and distribution of freshwater snails

Snail species of *Bulinus globosus, B. forskalii, Biomphalaria pfeifferi, Lymnaea natalensis, Pila ovala*, and *Lanistes lybicu*s were encountered in freshwater habitats classified into streams, ponds, and dams in Katsina-Ala Local Government Area. Similar snail species occurrence has been reported by Omudu and Iyough (2017) around the minor tributaries of the River Benue, Makurdi. The high frequency of occurrence of snail species in the pond may be because of habitat preference and the abundance of the pond. The ponds constituted 34.29% of the total water bodies surveyed. *B. globosus* had the highest numerical abundance, with 7,302 snails found in all 3 habitat types. The wide distribution of these species may be attributed to their ability to tolerate wide ranges of ecological factors (Asor and Arene, 2010). The occurrence of snail species throughout the year in some habitats, like ponds, could be a result of the permanent nature of the water bodies.

Human water contact activities seem to have an influence on snail occurrence and distribution. In part of the community where snails were found throughout the year, there was brick making, vegetable farming, and fishing. The banks of the habitats were also littered with human excrement as a result of the long stay of inhabitants in the area. This finding agrees with the findings of several authors that the population of schistosome intermediate hosts is often associated with moderate organic pollution (Houmsou, 2011; Omudu and Iyough, 2017). Sam-Wobo *et al*. (1991) suggested that snails in polluted water may use faeces as food or may benefit from the general eutrophication of the environment as a result of waste from the activities.

### (ii) Abundance of fresh water snails and their relationship to rainfall

*B. globosus* was the most abundant snail species encountered in this study, accounting for 29.43 % of the total snail species collected. This is similar to the findings of Omudu and Iyough (2017) and Asor and Arene (2010). Marked changes or fluctuations in the abundance of snail species in the freshwater habitats of Katsina-Ala seemed to be influenced by rainfall. In the stream habitats, there was a drastic drop in the population of *B. forskalii*. This may be because heavy rainfall increases the water level and the flood, resulting in the washing away of this snail species. This disagrees with the findings of Omudu and Odeh (2004), who stated that *B. forskalii* flourishes under flood conditions and their population seems to undergo rapid increase. The steady increase in population of *B. globosus* in ponds during the rainy season confirms the effect of rainfall on snails’ abundance. This steady increase was because these habitats were not flooded. The high numerical abundance of snails in the rainy season than in dry season also confirms the effects of rainfall on snails’ abundance.

Fluctuations in the population density of snails may not be dependent only on climate or general seasonal ecological conditions, but often significantly on peculiar local conditions caused by various factors or by the interference of man in those habitats.

### (iii) Examination of the snail for schistosome infection

It was established from the cercaria shedding experiment carried out that *B. globosus, B. forskalii, and B. pfefferi* are the vectors of schistosomiasis in Katsina-Ala, Benue State, Nigeria. This agrees with the reports of Asor and Arene (2010), who stated that *B. globosus, B. forskalii*, and *B. pfeifferi* were the vectors of schistosomiasis in the urban city of Port Harcourt and the Odau community of Rivers State, Nigeria. It was, however, different from the findings of Obonyo *et al*. (2010), who reported that *B. forskalii* was the vector of *S. intercalatum* in the urban city of Port Harcourt. The role of these snails as vectors of *S. haematobium* has been reported in other parts of the world (Christensen and Frandsen, 1985).

## 5.0 Conclusion

Based on the findings of this present research, the study calls for consideration of snail control to be integrated with PMDA campaigns to achieve transmission interruption and also recommends the strengthening of the existing schistosomiasis desk in the country through the employment of qualified personnel to handle the data collation, entry, and analysis.

## Supporting information

Ethical Clearance

## AUTHORS’ CONTRIBUTIONS

**James Agada Okete: C**onceptualized, designed, and managed the data of the study.

**Eme Effiong Etta:** Undertook the fieldwork and collection of data.

**Patricia Edodi Ikika:** Performed data analysis and interpretation.

**Theophilus Effiong Akpe:** Carried out data visualization, validation, and supervision.

**Benedict Onu Onoja:** Prepared the original manuscript, edited and submitted it (Corresponding Author).

## AUTHOR APPROVALS

All authors contributed to the development of the final manuscript and approved its submission. Also, this manuscript has not been accepted or published anywhere.

## APPENDIX I

### LIST OF GLOSSARIES

1. **Bilharziasis**: Another name for **schistosomiasis**, a parasitic disease caused by blood flukes (schistosomes) that live in certain types of freshwater snails before infecting humans.
2. **Biomphalaria pfeifferi**: A species of freshwater snail that acts as an intermediate host for **Schistosoma mansoni**, which causes intestinal schistosomiasis.
3. **Brevifurcate cercariae**: A type of larval stage of trematode parasites (including schistosomes) characterized by a short tail with forked ends. This stage actively swims in water to infect humans.
4. **Bulinus forskalii**: A species of freshwater snail that serves as an intermediate host for **Schistosoma haematobium**, which causes urogenital schistosomiasis.
5. **Bulinus globosus**: Another freshwater snail species that also transmits **Schistosoma haematobium**. It is a key vector in many parts of sub-Saharan Africa.
6. **Cercariae**: The free-swimming larval stage of schistosome parasites that emerge from infected snails and can penetrate human skin, initiating infection.
7. **Epibiotic**: Refers to an organism that lives on the surface of another living organism (host) without harming it. For example, algae growing on snails.
8. **Lanistes lybicus**: A species of large freshwater snail found in African water bodies. It is **not** typically involved in schistosomiasis transmission.
9. **Lymnaea natalensis**: A species of freshwater snail known to be involved in transmitting **liver flukes**, but **not schistosomes**.
10. **Pila ovata**: A large freshwater snail found in Africa, not known to be involved in schistosomiasis transmission but may serve other ecological roles.
11. **Praziquantel**: An antiparasitic medication used to treat **schistosomiasis** and other fluke infections. It is safe, effective, and commonly used in mass treatment programs.
12. **Praziquantel Mass Drug Advocacy**: Efforts to promote and support widespread administration of **praziquantel** to populations at risk of schistosomiasis, including public education and policy support.
13. **Schistosomiasis**: A **parasitic disease** caused by **Schistosoma** worms. It affects the intestines or urinary tract and is transmitted through contact with freshwater contaminated with cercariae.
14. **Schistosomes**: Parasitic flatworms (trematodes) of the genus **Schistosoma**. They infect humans via freshwater and are responsible for causing schistosomiasis.

## APPENDIX II

## LIST OF ABBREVIATION

FMOH: Federal Ministry of Health
MDA: Mass Drug Administration
NGOs: Non-Governmental Organization
NTDs: Neglected Tropical Diseases
PMDA: Praziquantel Mass Drug Administration
PZQ: Praziquantel
WHA: World Health Assembly

