## Supplementary figures and images for "Occurrence of snail intermediate host of schistosomiasis: implications for schistosomiasis control based on mass drug administration of praziquantel in Benue State, Nigeria"

### Ethical Clearance

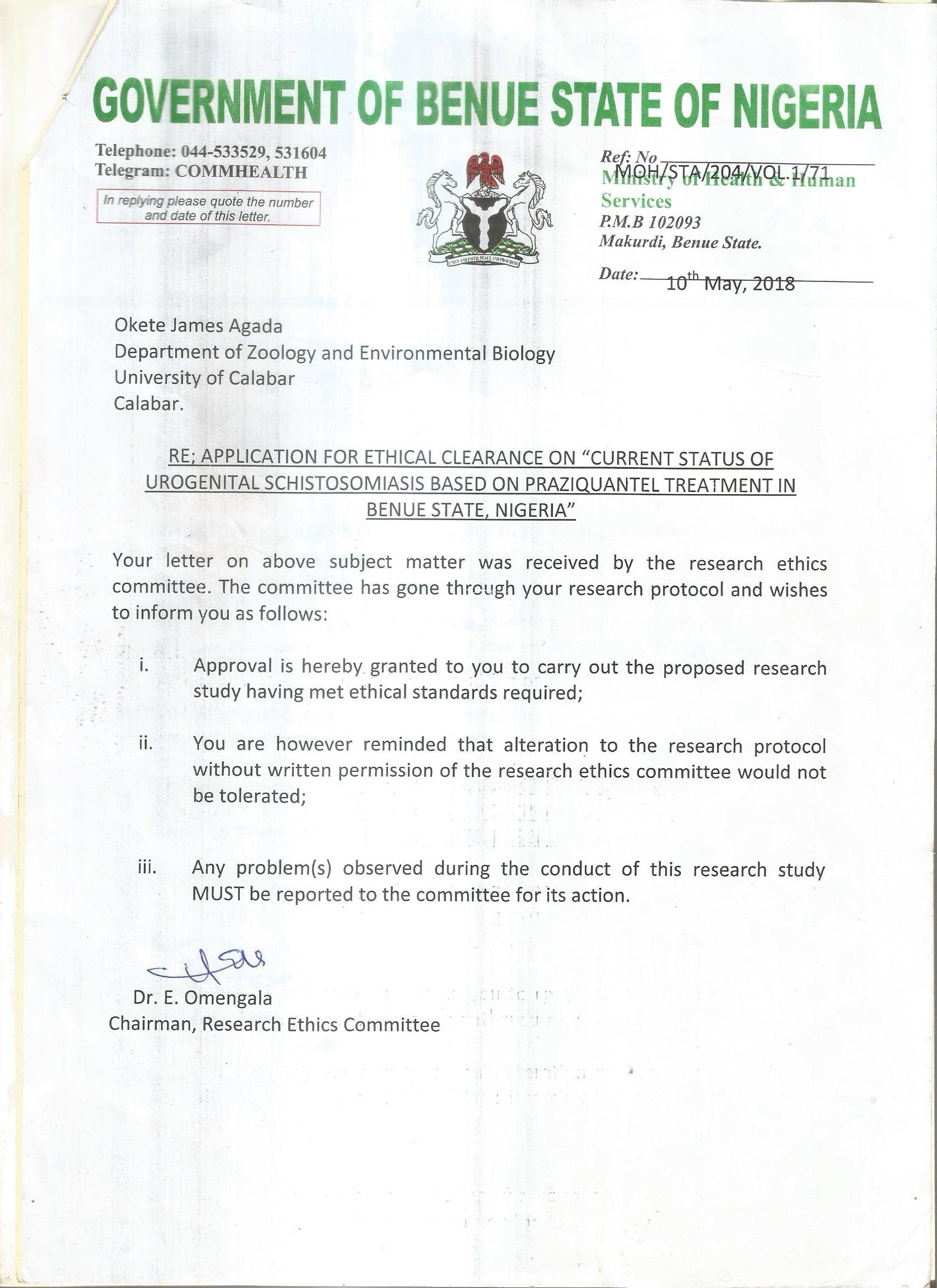
